# Cloning and characterization of Osmotin (*OsmSt*) alleles from potato cultivar ‘Kufri Chipsona 1’

**DOI:** 10.1101/2020.01.03.894030

**Authors:** Amanpreet Kaur, Anil Kumar

## Abstract

The present study was focussed to clone and sequence characterise alleles of osmotin from cDNA of *Solanum tuberosum* L. cultivar ‘Kufri Chipsona 1’. The genes vary in sizes as well as were found to have eleven point mutations throughout the coding sequence. One deletion of 7 bp was also found in smaller form of the gene having molecular weight of 19.91 KDa. Osmotin gene is known to impart resistance/tolerance to various fungal diseases in addition to its role as an osmoprotectant, thus, cloned osmotin alleles from important processing grade potato cultivar could become potential candidates for molecular breeding of potato.

## Introduction

Under the natural conditions, plants are continuously subjected to various biotic and abiotic stresses causing impaired cellular growth (Kumar et al. 2015). To sustain these stresses, plants have evolved a cascade of defense strategies involving release of various proteins such as pathogenesis related (PR) proteins (Li et al. 2015). On the basis of their biochemical properties and mechanism of action, PR proteins are divided into 17 families including β-1,3 glucanases (PR-2), chitinases (PR-3, 4, 8 and 11), thaumatin-like proteins (PR-5), ribosome inactivating proteins (PR-10), defensins (PR-12), thionins (PR-13) and lipid transfer proteins (PR-14) (Kaur et al. 2017a). Among these, PR-5 protein family is known for its antimicrobial activity, osmoregulation and role in plant development (Mani et al. 2012). PR-5 family is named as thaumatin-like protein (TLPs) due to its close similarity with sweet tasting protein thaumatin of plant *Thaumatococcus danielli* (Velazhahan et al. 1999). PR-5 proteins are classified into three types (acidic, neutral and basic) based on their isoelectric points (Kumar et al. 2015). Out of all the known PR-5 proteins, osmotin or osmotin-like proteins (OLPs) are well known for their role as osmoprotectants and anti-fungal agents (Chowdhury et al. 2015, 2017). They are reported to get accumulated under stress conditions leading to ion/solute compartmentalization and anti-phytopathogen activities (Kumar et al. 2015).

Osmotin, a multifacet stress inducible protein was firstly isolated from tobacco cells exposed to saline conditions (Choi et al. 2013). The protein have been cloned from various plants such as *Nicotiana tabaccum* (Lui et al. 1994, Evers et al. 1999, Zuker et al. 2001, Ouyang et al. 2005, Qin et al. 2006, Noori and Sokhansanj 2008, Parkhi et al. 2009, Goel et al. 2010, Rao et al. 2011, Husaini et al. 2012, Subramanyam et al. 2011), *Solanum nigrum* (Vasavirama and Kirti 2012), *Capsicum annuum* (Choi et al. 2013), *Solanum tuberosum* (Rivero et al. 2012), etc. The over-expression of osmotin have been reported to exhibit anti-fungal activity against diverse range of pathogens such as *Puccinia triticina* (Li et al. 2015), *Fusarium graminearum* (Datta et al. 1999, Mackintosh et al. 2007), *Phytophthora capsici, Fusarium oxysporum* (Zuker et al. 2001, Mani et al. 2012), *Phytophthora infestans* (Liu et al. 1994, 1996, Li et al. 1999), *Rhizoctonia solani* (Rivero et al. 2012), etc. The anti-fungal property of the protein attributes to its capability to bind to domains present on the fungal plasma membrane (Thevissen et al. 2005) which leads to dissipation of membrane integrity and can also inhibit spore germination through spore lysis (Koiwa et al. 1997). Osmotin has been reported to be induced in plants exposed to salt stress (Qureshi et al. 2007), cold stress (Patade et al. 2012), ethylene, wounding (Zhu et al. 1995) and biotic stresses (Elvira et al. 2008, Mukherjee et al. 2010, Choi et al. 2013). Due to its anti-fungal properties and highly stable nature, osmotin is now considered as a natural food preservative (Thery et al. 2019). Osmotin also shows structural and functional similarity to insulin-sensitizing hormone, adinopectin, which further highlights the importance of protein in therapeutics (Kumar et al. 2015).

In the present study, we report cloning and characterization of two osmotin alleles from an important processing grade Indian potato cultivar ‘Kufri Chipsona 1’. The attempts were also made to highlight characteristic features and structure of osmotin proteins encoded by the cloned alleles using bioinformatics tools. The cloned gene can be used in molecular breeding of potato and other plants to improve resistance/ tolerance against various abiotic and biotic stress conditions.

## Material and Methods

### Plant material and chemicals

Microshoot cultures of *Solanum tuberosum* L. cultivar ‘Kufri Chipsona 1’ (CS-1) maintained at plant tissue culture laboratory, Thapar Institute of Engineering & Technology, Patiala through regular subculture on MS1 [basal Murashige and Skoog (1962) medium containing 10μM silver nitrate] medium (Kaur et al. 2017b) were used for cloning of osmotin (*OsmSt*) gene. The cultures were incubated at 25±2 °C, under photoperiod duration of 16/8 h day/night and photo intensity of 50 μmol.m^2^/s provided by cool fluorescent lamps (Philips Ltd., India). All tissue culture grade chemicals and enzymes were purchased from HiMedia Laboratories Ltd. (Mumbai, India) and ThermoScientific Laboratories Ltd. (Mumbai, India) respectively.

### Cloning of *OsmSt* gene

Total RNA was isolated from the microshoots of potato cultivar CS-1 following CTAB method (Chang et al. 1993) and cDNA was synthesized using RevertAid First Strand cDNA synthesis kit (Fermentas Life Sciences, U.S.A) according to manufacturer’s guidelines. cDNA diluted by 100 times was used as template for gene amplification using primers [Pair 1: Forward primer 5’-CGGGATCCCGCCACAAACATGGCCTACTT-3’, Reverse Primer 5’-CGAGCTCGTTACTTAGCCTCTTCATCACTTGAAG-3’, Pair 2: Forward Primer 5’-CGGGATCCGCCAGACGTGGGTCATAAAT-3’, RP 5’-CGAGCTCGCGGGCCAGTAACACCATTAG-3’] having overhangs with restriction site for BamHI and SacI. The PCR amplification was carried out in GenAmp 2700 thermocycler (Applied Biosystem, USA) using reaction mixture consisting of template (cDNA), Taq Polymerase (1 U), dNTPs mixture (100 μmol), 2 μl reaction buffer (10X), primers (10 nmol each) and sterile Milli-Q water (Millipore India, Bangalore) to make up the volume upto 20 μl. The amplification conditions includes initial denaturation at 94 °C for 5 min followed by 31 cycles of 94 °C (1 min), 60 °C (45 sec), 72 °C (1.5 min) and final extension at 72 °C for 5 min. The amplified DNA fragments separated on ethidium bromide stained agarose gel (1% w/v) were visualised on U.V. transilluminator and documented using GelDoc system (Biosystematica, U.S.A).

PCR products was gel purified using the PCR purification kit (Qiagen, USA) and restricted using BamHI and Sac1 for directional cloning into pBI121 binary vector (Chen et al. 2003). The ligated products were introduced into *E. coli* strain DH5α and propagated on Luria Agar plates containing kanamycin (50 mg/l) as selection antibiotic. The modified plasmids were isolated from the selected clones and sequenced in Applied Biosystems automatic sequencer (DNA Sequencing Facility, Department of Biochemistry, South Campus, Delhi University, New Delhi, India) using gene specific primers.

### Analysis of sequence data

The sequences of osmotin alleles (*OsmSt1* and *OsmSt2*) thus obtained were compared with reported nucleotide and protein sequences those available in GenBank/ EMBL databases using BLAST program (Altschul et al. 1997) accessible through NCBI (National Centre for Biotechnology Information – http://www.ncbi.nlm.nih.gov). Multiple sequence alignments were performed using MULTALIN program (http://multalin.toulouse.inra.fr/multalin/). The identification of the ORF and amino acid sequence coded by the cloned genes were deduced using the ORF finder program provided by the NCBI (http://www.ncbi.nlm.nih.gov/gorf/gorf.html). Homology searches for deduced amino acid sequence were performed with BLAST and multiple sequence alignments were carried out with OsmSt1 and OsmSt2 protein sequences using CLUSTALW programme available online at www.ebi.ac.uk. The alignment was modified to highlight conserved regions using GeneDoc programme (www.nrbsc.org/gfx/genedoc). Potential transmembrane segments were identified using TMHMM-2.0 (http://www.cbs.dtu.dk/services/TMHMM-2.0) programme. The glycosylation and phophorylation sites were identified using NetNGlyc 1.0 and NetPhos 2.0 Server from technical University of Denmark (http://genome.cbs.dtu.dk//cgi-bin/webface?jobid=netNglyc and http://genome.cbs.dtu.dk//cgi-bin/webface?jobid=netphos). The molecular mass and isoeletric point of both proteins were computed using Compute pI/Mw tool from ExPASy (Expert ProteinAnalysis System) bioinformatics resource portal (http://web.expasy.org/compute_pi/) (Bjellqvist et al. 1993). Similarly for calculating the amino acid composition of the predicted amino acid sequences, the ProtParam tool of ExPASy proteomics server of the Swiss Institute of Bioinformatics (SIB; http://expasy.org/tools/) was used. Domains and family of both proteins were identified using interproscan tool of European Bioinformatics Institute (http://www.ebi.ac.uk/Tools/pfa/iprscan/) (Quevillon et al. 2005).

### Molecular modelling and prediction of disulphide bridges

The amino acid sequences of OsmSt1 and OsmSt2 were first subjected to PSI-BLAST searches (Altschul et al. 1997) to find homologous template. The single highest scoring template providing maximum coverage with 100% confidence was used for modelling. Consequently, the structure of PR-5d of *N. tabaccum* was retrieved and used to develop model of OsmSt1 and OsmSt2 using Phre2 protein fold recognition server (www.sbg.bio.ic.ac.uk). The three dimensional structure was predicted through alignment of target sequence with template sequence.

The presence of disulphide bonds and state of cysteine residues in OsmSt1 and OsmSt2 were predicted using DiANNA 1.1 web server (clavius.bc.edu). The scores were produced for each pair of cysteine in the amino acid sequences and Edmonds-Gabov maximum weight matching algorithm was used to pair cysteine residues for formation of disulphide bonds (Ferre and Clote 2005).

## Result and Discussion

Osmotin and its various isoforms have been cloned from many plants (Capelli et al. 1997, Pla et al. 1998, Jami et al. 2007, Singh et al. 2017, Mani et al. 2012, Li et al. 2015). The presence of osmotin isoforms in same plant have been previously reported in *Piper colubrinum* (Mani et al. 2012). The two isoforms were reportedly distinct from each other in terms of size, pI value and molecular structure. In the present study, we report cloning and sequence characterization of osmotin alleles from an important Indian potato cultivar ‘Kufri Chipsona 1’ (CS-1).

cDNA synthesised using high quality, intact RNA (416.7 ng/μl; A_260/280_ 2.09) isolated from actively growing microshoots of CS-1 yielded amplification of 733bp for both *OsmSt1* and *OsmSt2* alleles. BLAST analysis of the sequences revealed high similarity of PCR products with the osmotin. More than 99% nucleotide sequence similarity was observed between osmotin alleles from CS-1 and other Solanum species [*S. tubersosum*; XP_006358887.1 (99.45%), *S. lycopersicum*; NP_001296216.1 (99.45%), *S. pennellii*; XP_015083272.1 (99.45%)]. It have been previously reported that osmotin gene is highly conserved among the Solanaceae family (Kumar et al. 2015) and slight variation in nucleotide level may be due to presence of single nucleotide polymorphism (SNPs) in the sequence (Jami et al. 2007). It has also been reported that GC content is used for hierarchical classification of a species (Wayne et al. 1987). The reported GC content for land plants ranges between 30–50% (Lynch 2007). In case of cloned *OsmSt1* and *OsmSt2* genes, GC content was found as 46.52% which fall in the reported range.

Alignment of *OsmSt1* with *OsmSt2* showed presence of single nucleotide polymorphism (at 418, 555, 651, 664, 683, 685, 688-692 positions of nucleotides) and a deletion between 674 to 681 bp (Figure 1). The open reading frame (ORF) of *OsmSt1* was found to have 561 nucleotides encoding 186 amino acids whereas *OsmSt2* was slightly larger having 576 nucleotides encoding 192 amino acids. Although showing high amino acid sequence similarity, the functional properties were found to vary between the OsmSt1 and OsmSt2 proteins due to presence of additional six amino acids in OsmSt2. The calculated molecular mass of OsmSt1 was found to be 19.91 kDa with an isoelectric point (pI) of 4.81, whereas OsmSt2 was having molecular mass of 20.55 KDa and was slightly acidic (pI 6.06).

**Figure 1:**
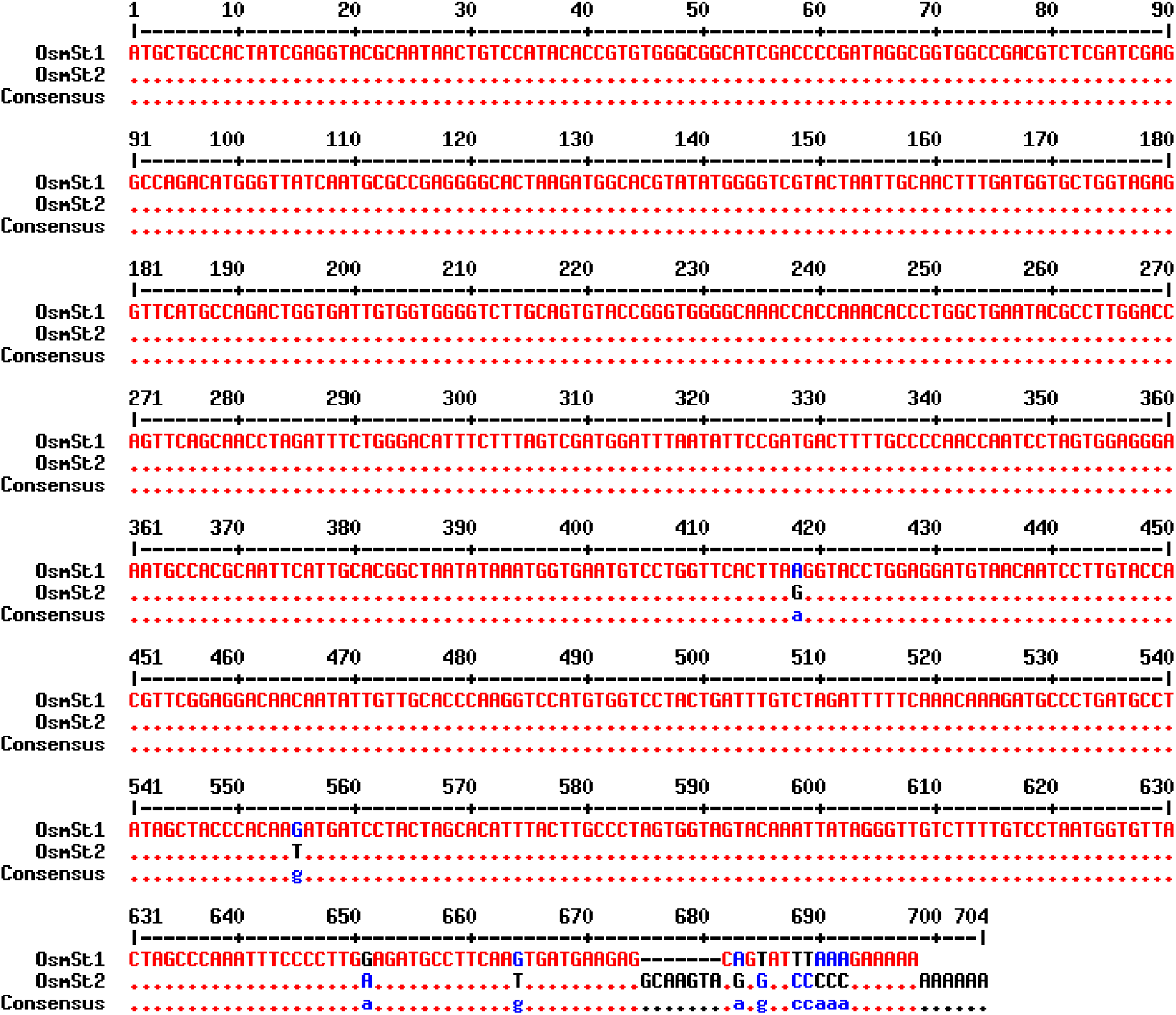
Sequence alignment of osmotin alleles cloned from potato cultivar ‘Kufri Chipsona 1’ showing point mutations (at 418^th^, 555^th^, 651^st^, 664^th^, 683^rd^, 685^th^, 688-692^nd^ positions) and deletion between 674th and 681st position in coding sequence of Osmotin allele 1 (*OsmSt1*)

PSI-BLAST with 1 iteration revealed high similarity of CS-1 osmotin with other species (Table 1, Figure 2). The Osmotin protein from these species is reported to play a role in plant defense system against abiotic stress and fungal invasions (Castillo et al. 2005, Kumar et al. 2015). Therefore, there is a strong possibility that OsmSt1 and OsmSt2 from CS-1 is likely to show similar antifungal activities.

**Table 1:** BLAST analysis of amino acid sequences

| Accession number | Description | Amino acid similarity (%) | E-Value |
| --- | --- | --- | --- |
| XP_006358887.1 | Osmotin-like protein from *Solanum tuberosum* | 99.45 | 5e-131 |
| NP_001296216.1 | Osmotin-like protein (TPM-1) from *Solanum lycopersicum* | 99.45 | 7e-131 |
| XP_015083272.1 | Osmotin-like protein (OSML13) from *Solanum pennellii* | 99.45 | 1e-130 |
| AAU93853.1 | Osmotin-like protein (A13) from *Solanum phureja* | 98.90 | 1e-129 |
| P50701.1 | Osmotin-like protein (OSML13) from *Solanum commersonii* | 98.90 | 1e-130 |
| QC092097 | Osmotin from *Solanum tuberosum* | 97.84 | 3e-131 |
| AAS48588.1 | Putative Osmotin-like protein from *Brassica juncea* | 93.44 | 2e-124 |
| AAL87640.1 | Osmotin-like protein from *Solanum nigrum* | 93.96 | 2e-124 |
| NP_001311827.1 | Osmotin-like protein from *Capsicum annuum* | 93.89 | 2e-122 |
| CAA43854.1 | Osmotin from *Nicotiana tabacum* | 87.85 | 2e-113 |
| CAA61411.1 | Osmotin from *Arabidopsis thaliana* | 70.43 | 2e-84 |
| ABX71220.1 | Osmotin from *Piper colubrinum* | 75.46 | 3e-80 |

**Figure 2:**
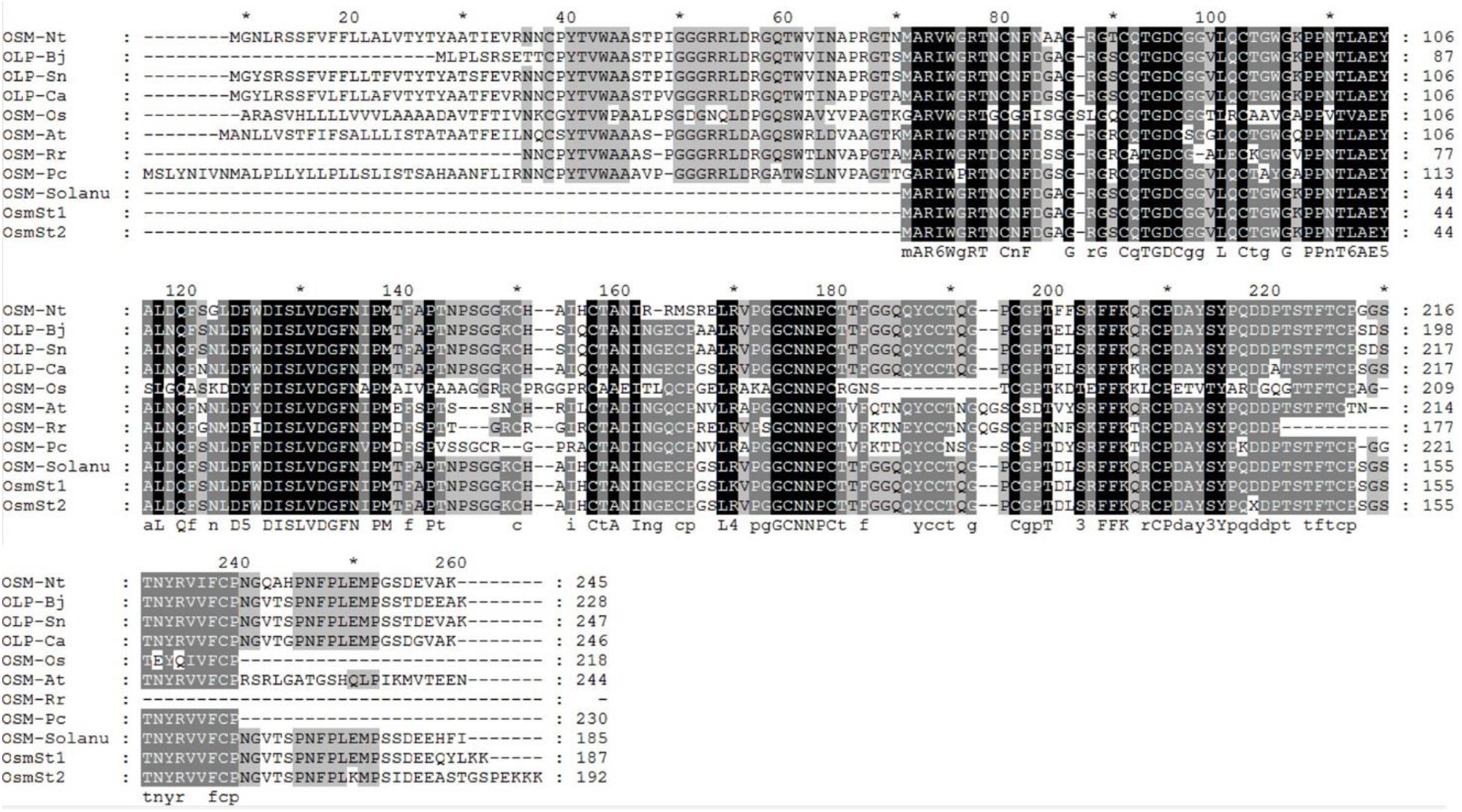
Multiple sequence alignment of amino acid sequences of ‘Kufri Chipsona 1’ osmotin (OsmSt1 and OsmSt2) with osmotin (OSM) and osmotin-Like-proteins (OLP) from different taxas viz. *Nicotiana tabaccum* (OSM-Nt, CAA43854.1), *Brassica juncea* (OLP-Bj, AAS48588.1), *Solanum nigrum* (OLP-Sn, AAL87640.1), *Capsicum annuum* (OLP-Ca, NP_001311827.1), *Oryza sativa* (OSM-Os, AAB67852.1), *Arabidopsis thaliana* (OSM-At, CAA61411.1), *Rosa roxburghii* (OSM-Rr, ABC39615.1), *Piper colubrinum* (OSM-Pc, ABX71220.1), *Solanum tuberosum* (OSM-Solanu, QC092097.1). Black coloured highlighted sequences indicates the conserved regions of the protein.

Both proteins were found to be acidic in nature with presence of more acidic amino acids in comparison with basic amino acids (https://web.expasy.org/protparam/). It was also found that some amino acids such as Gly (11.9-12.0%), Pro (9.7-9.9%), Ser (7.0-7.3%), Thr (8.6-8.9%), Cys (7.8-8.1%) and Asn (6.8-7.0 %) occurred more frequently whereas other amino acids such as Ala (4.3-4.7%), Arg (3.6-3.8%), Asp (5.2-5.9%), Gln (4.2-4.3%), Glu (2.6-2.7%), His (1.0-1.6%), Leu (4.2-4.3%), Met (1.6%), Lys (1.6-3.6%) and Trp (1.6%) occurs less frequently. It has been previously reported that overall amino acid composition of the protein have an important role in evolution (King and Jukes 1969). Further, the quality and structure of protein is also found to be dependent upon amino acid composition (Dyer 1971). Osmotin protein is reported to be rich in cysteine residues scattered all over the sequence. These residues form disulphide bridges which is a signature of PR-5 family proteins (Min et al. 2004, Jami et al. 2007). Generally, the presence of 16 cysteine residues have been reported in osmotin (Kumar et al. 2015), but in the present study, only 15 cysteine residues were present which lead to formation of 7 disulphide bridge only (Figure 3). The presence of disulphide bridges is known to provide high stability to the protein (Mani et al. 2012). The instability index (II) calculated for OsmSt1 and OsmSt2 using ProtPram tool of Expasy server were found to be about 38.90 which indicates that the proteins are highly stable (https://web.expasy.org/protparam/).

**Figure 3:**
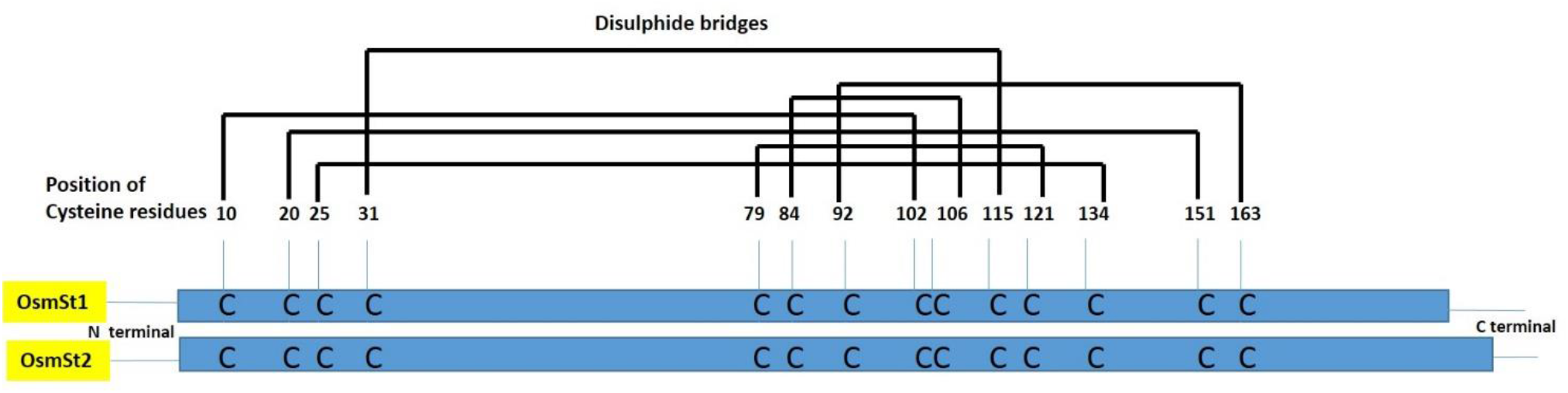
Prediction of disulphide bridges between cysteine residues of proteins (OsmSt1 and OsmSt2) encoded by osmotin alleles cloned from potato cultivar ‘Kufri Chipsona 1’

Homology models of OsmSt1 and OsmSt2 are shown in Figure 4. The structures are typical to PR-5 family proteins with three distinct domains. OsmSt1 and OsmSt2 were found to be similar to each other having nine *β*-sheets and four *α*-helices (Figure 4a-b). The models were produced using PR-5d protein from *N. tabacum* (d1auna_b.25.1.1; Tax ID 4097) as template which provide 89% and 86% coverage (166 amino acids) to OsmSt1 and OsmSt2 repectively with 100% confidence. The model of template was also produced (Figure 4c) which was also found to have three distinct domains with 12 *β*-sheets and four *α*-helices. Our predicted models are in line with the structural information of PR-5 family proteins (zaematin, thaumatin-like protein, Osmotin) previously reported (Mani et al. 2012).

**Figure 4:**
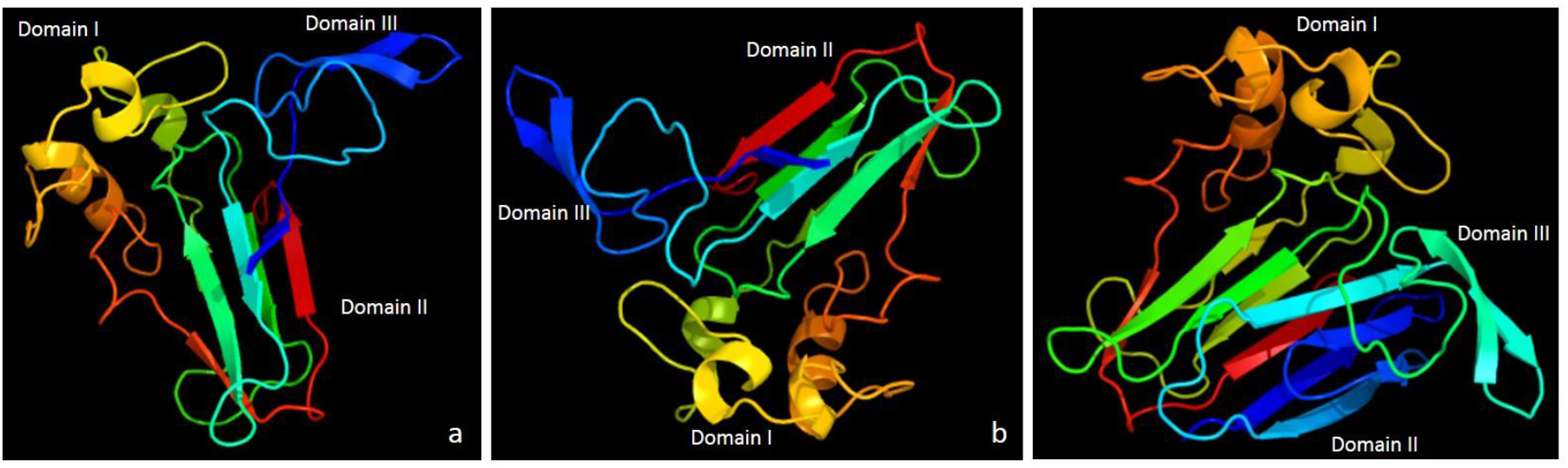
Molecular modelling of Osmotin using PR-5 protein from *N. tabaccum* (TaxID:4097) as template. a-b) homology models of OsmSt1 and OsmSt2 respectively showing three domains of protein consisting of nine β-sheets and four α-helices connected by various loops c) homology model of template showing typical structure of thaumatin-like-protein consisting of twelve β-sheets and four α-helices.

Glycosylation and phosphorylation are two important post-translational modification events which modify protein properties by altering its distribution, folding and stability (Ciesla et al. 2011, Haltiwanger and Lowe 2004). Thus, in the present study, presence of glycosylation and phosphorylation sites were analysed using NetPhos 2.0 and NetGlyc 1.0 servers. OsmSt1 and OsmSt2 from CS-1 do not show the presence of glycosylation sites whereas the presence of 16 phosphorylation sites were observed (Figure 5). Similar findings were observed from reported Osmotin*-like* protein of *Solanum tuberosum* accession XM_006358825.2. Out of 16 detected phosphorylation sites, 11 were at serine residue, 4 at threonine residue and 1 at tyrosine residue (Figure 5). These results were in line with previous findings which suggested the occurance of phosphorylation on serine residues as most common phenomena (Thomason and Kay 2000). It has also been reported that during interaction with fungal pathogen, osmotin utilizes signal transduction pathway which is dependent on phosphorylation and dephosphorylation events (Kim et al. 2002).

**Figure 5:**
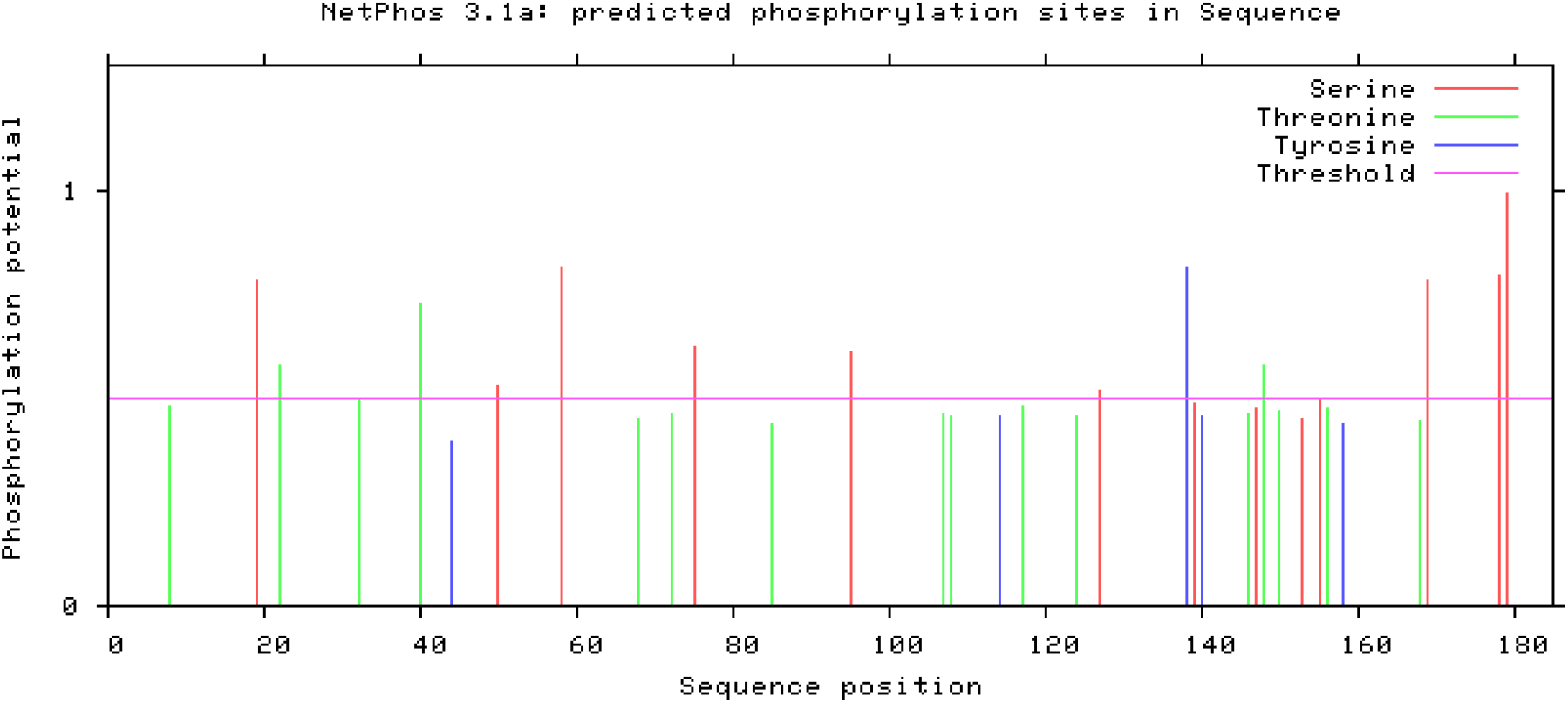
Prediction of phosphorylation sites in osmotin proteins (OsmSt1 and OsmSt2) using NetPhos 2.0 server (http://genome.cbs.dtu.dk//cgi-bin/webface?jobid=netphos)

In conclusion, alleles of osmotin were cloned and characterized from *Solanum tuberosum* cultivar ‘Kufri Chipsona 1’. The proteins encoded (OsmSt1 and OsmSt2) by these alleles were found to be highly similar but posses different physio-chemical properties (pI and molecular weight) which could determine their mode of action. The structural analysis of proteins revealed presence of three distinct domains and various disulphide bridges which is a characteristic feature of osmotin. The cloned alleles from CS-1 could be a candidate for in-depth functional and evolutionary studies and could lead to potential molecular breeding programmes for generation of disease resistant/ salt tolerant potato varieties.

## Acknowledgement

Amanpreet Kaur is thankful to University Grants Commission (UGC), New Delhi for the award of Maulana Azad National Fellowship for minority students. TIFAC-CORE is thanked for the facilities to carry out research work.

## Author contribution

Amanpreet Kaur conducted the experiments, compiled data, analyzed the results and wrote the initial draft of the manuscript, Amanpreet Kaur and Anil Kumar conceived and designed the experiments, Anil Kumar finalized the manuscript.

